# Characterization of membrane vesicles in *Alteromonas macleodii* indicates potential roles in their copiotrophic lifestyle

**DOI:** 10.1101/2022.09.27.509651

**Authors:** Eduard Fadeev, Cécile Carpaneto Bastos, Jennifer Hennenfeind, Steven J Biller, Daniel Sher, Matthias Wietz, Gerhard J Herndl

**Affiliations:** Department of Functional and Evolutionary Ecology, Bio-Oceanography and Marine Biology Unit, University of Vienna, Vienna, Austria; Department of Biological Sciences, Wellesley College, Wellesley, USA; Department of Marine Biology, Leon H. Charney School of Marine Sciences, University of Haifa, Haifa, Israel; Deep-Sea Ecology and Technology, Alfred Wegener Institute Helmholtz Center for Polar and Marine Research, Bremerhaven, Germany; Max-Planck Institute for Marine Microbiology, Bremen, Germany; NIOZ, Department of Marine Microbiology and Biogeochemistry, Royal Netherlands Institute for Sea Research, Utrecht University, Den Burg, The Netherlands

**Keywords:** bacterioplankton, EVs, transporters, extracellular enzymes, iron uptake, moonlighting proteins

## Abstract

Bacterial membrane vesicles (MVs) are likely abundant in the oceans. Based on observations from non-marine bacteria, MVs are involved in a range of physiological processes and play important roles in interactions between microbial cells. In this study we characterized MV production of six different strains of *Alteromonas macleodii*, a cosmopolitan marine bacterium. *A. macleodii* strains produced MVs at rates of up to 30 MVs cell^-1^ generation^-1^. The produced MVs had high morphological diversity that could potentially define their functional roles. Proteomic characterization revealed that MVs are rich in membrane proteins related to iron and phosphate uptake, as well as proteins with potential functions in biofilm formation. Furthermore, MVs were harboring hydrolytic enzymes. Taken together, our results suggest that in the largely oligotrophic oceans, *A. macleodii* MVs may support its growth through generation of extracellular “hotspots” that facilitate access to essential substrates. This study provides an important basis for further investigation of the ecological relevance of MVs in heterotrophic marine bacteria.

## Introduction

Production of extracellular vesicles (EVs) occurs in all three domains of life (Deatherage and Cookson 2012). Once considered as cell debris, it is now realized that EVs play an important role in microbial physiology and ecology. Both gram-negative and -positive bacteria produce membrane vesicles (MVs; Nagakubo, Nomura and Toyofuku 2019). The MVs are lipid structures, range between 20-400 nm in diameter, and carry various biomolecules including proteins, enzymes, and nucleic acids (Nagakubo, Nomura and Toyofuku 2019). The most common mechanism of MV release in gram-negative bacteria is through blebbing of the outer membrane, releasing so-called outer-membrane vesicles (Schwechheimer, Sullivan and Kuehn 2013). This type of MVs is typically of spherical structure with an external lipopolysaccharide and an inner phospholipid layer that encloses fragments of the periplasmic matrix (Deatherage *et al*. 2009). Several other types of MVs were recently characterized in gram-negative bacteria, such as outer-inner membrane vesicles that contain both layers of the cellular membrane and have cytoplasmic content (Pérez-Cruz *et al*. 2015). Furthermore, tube-shaped membranous structures can represent a chain of MVs or enclosed extensions of cellular outer membranes (Fischer *et al*. 2019). The exact mechanisms behind the production of different types of MVs are still not fully understood (Toyofuku, Nomura and Eberl 2019). Although it has been suggested that different types of MVs might be associated with different biological functions, here we will address them all as MVs.

Most knowledge on the functional roles of MVs originates from research on bacteria-host interactions (Nagakubo, Nomura and Toyofuku 2019). For example, human pathogenesis studies revealed that MVs can deliver virulence and toxin cargoes and lead to a range of serious illnesses, ranging from lung infection to cancer (Caruana and Walper 2020; Chronopoulos and Kalluri 2020). However, MVs also play key roles in bacteria-bacteria interactions, such as quorum sensing and microbial “warfare” in *Pseudomonas aeruginosa* (Tashiro *et al*. 2013; Lin *et al*. 2018), as well as distribution of antibiotic resistance genes between bacterial lineages (Yaron *et al*. 2000; Lee *et al*. 2013). Furthermore, MV production can be stimulated by environmental triggers (Orench-Rivera and Kuehn 2016) such as oxygen availability (Toyofuku *et al*. 2014) or iron starvation (Roier *et al*. 2016). This body of evidence from other microbial systems suggests that MVs likely also have important functional roles in marine microbial communities (Schatz and Vardi 2018).

A recent exocellular DNA study suggested that most, if not all, pelagic marine bacteria produce MVs (Linney *et al*. 2022). However, to date, MV production has only been shown for few marine bacterial isolates (Biller *et al*. 2022). In two isolates of *Prochlorococcus*, the most abundant marine cyanobacterium, a continuous production of MVs harboring a diverse array of cellular compounds has been observed (Biller *et al*. 2014, 2017). Some *Prochlorococcus* MVs contain heterogeneous fragments of DNA, which might be associated with horizontal gene transfer (recently termed “vesiduction”; Soler and Forterre 2020). Other *Prochlorococcus* MVs are characterized by a highly heterogeneous composition of nucleic acids, proteins, and secondary metabolites, the functional roles of these are yet to be identified (Biller *et al*. 2021). Among the heterotrophic members of marine microbial communities, MV production was observed in *Shewanella* (Gammaproteobacteria; Frias *et al*. 2010) and *Formosa* spp. (Bacteroidia; Fischer *et al*. 2019). In both isolates, the produced MVs had a high content of membrane and periplasmic proteins associated with organic matter and nutrient processing, with potential ecological importance.

To further elucidate the functional roles of MVs in heterotrophic marine bacteria, we investigated MV production and characterized MVs in the marine bacterium *Alteromonas macleodii*. This gammaproteobacterial genus is a prominent member of pelagic bacterial communities in the ocean (García-Martínez *et al*. 2002), and can be associated with various phytoplankton groups (Kearney *et al*. 2021), such as *Prochlorococcus* (Biller, Coe and Chisholm 2016). *A. macleodii* occurs both free-living and surface attached (i.e., in biofilms) (Ivars-Martinez *et al*. 2008), and has been extensively studied through multiple genome-sequenced isolates from various oceanic regions (López-Pérez *et al*. 2012, 2013; Ivanova *et al*. 2015). This genus harbors a high degree of genomic flexibility through intra- and inter-species exchange of genomic islands and plasmids (López-Pérez, Gonzaga and Rodriguez-Valera 2013; Fadeev *et al*. 2016; López-Pérez, Ramon-Marco and Rodriguez-Valera 2017), facilitating ecological differentiations and niche specialization (López-Pérez and Rodriguez-Valera 2016; Koch *et al*. 2020). Nevertheless, all *A. macleodii* strains known to date are characterized as opportunistic copiotrophs, for instance specialized in polysaccharide degradation (Neumann *et al*. 2015; Koch *et al*. 2019) and rapid uptake of organic carbon (Pedler, Aluwihare and Azam 2014), as well as scavenging of iron through the production of siderophores (Manck *et al*. 2022). We hypothesize that some of these physiological and ecological traits of *A. macleodii* may involve MVs.

In this study we investigated the production and functional potential of MVs in *A. macleodii*. We determined the MVs production rates of six different *A. macleodii* strains and compared the morphologically variable MVs populations. Supported by protein characterization of isolated MVs, we propose various potential roles of MVs in biological and biogeochemical traits of *A. macleodii*, which might contribute to the cosmopolitan occurrence of this marine bacterium.

## Results and Discussion

### *A. macleodii* produces morphologically diverse MVs

The production of MVs has been previously investigated in isolates of several pelagic marine bacteria, such as *Prochlorococcus* (Biller *et al*. 2014, 2021), *Pelagibacter* (SAR11; Biller *et al*. 2022), and *Shewanella* (Frias *et al*. 2010; Baeza and Mercade 2021). We extended these observations by characterizing MVs in six strains of the cosmopolitan marine bacterium *A. macleodii* exhibiting high morphological diversity. Using cryogenic electron microscopy (cryo-EM), heterogeneous MVs in the size range of 40-300 nm were observed (Figure 1). Further investigation of the MVs using nanoparticle tracking analysis (NTA) revealed a bi-modal size distribution with a pronounced main population of MVs with a diameter of 80-120 nm and an additional population with a diameter of 200-240 nm (Figure 2). This size range of *A. macleodii* MVs is overall in agreement with observations in other marine bacteria (Biller *et al*. 2014, 2017, 2022). Microscopic images also revealed some larger MVs of > 300 nm, most of which were probably removed during the purification process that included. Thus, observations of these larger MVs suggests that the size cut-off of < 0.22 μm applied here (and in previous studies of MVs in marine bacteria) could potentially skew the observations towards sub-populations of smaller spherical MVs.

**Figure 1.**
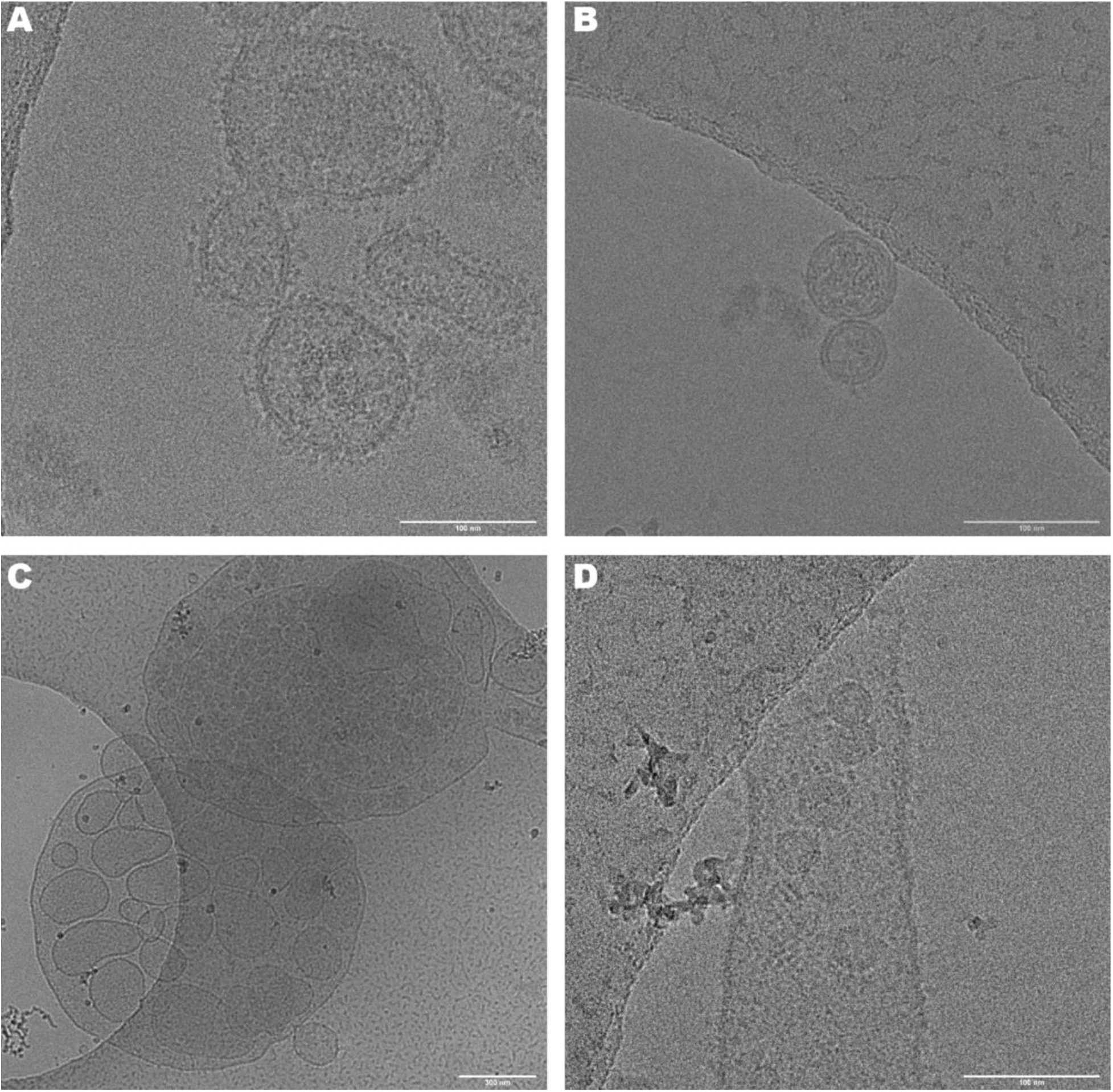
Distinct morphological shapes in *A. macleodii* MVs, visualized using cryo-EM. (**A**) strain BS11 - protrusions of potentially lipopolysaccharide or proteins surrounding the MVs. (**B**) strain HOT1A3 - outer inner-MVs. (**C**) strain BGP6 - micron-large membrane-bound structures. (**D**) strain BS11 - tube-shaped membranous structures. Note the larger scale bars in panels C and D.

**Figure 2.**
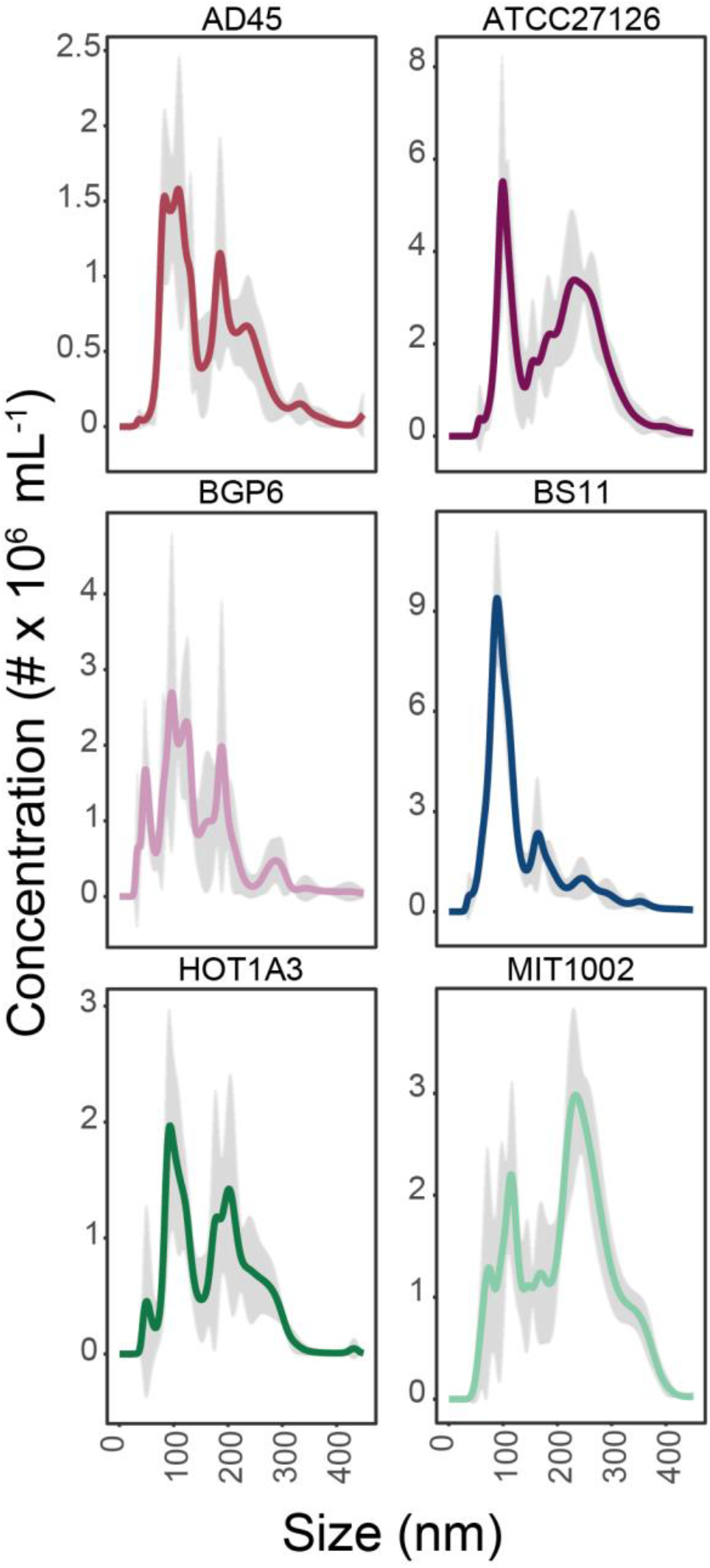
Size characterization of MVs in *A. macleodii* strains. Each plot represents the NTA size distribution of MVs; the shaded area representing the standard deviation between the technical replicates of the NTA runs.

Previous estimates of MV production rates in pelagic marine bacteria range from few to several dozen MVs released per bacterial cell (Biller *et al*. 2014, 2022; Kwon *et al*. 2019). In the six strains of *A. macleodii*, we observed MV production rates, calculated based on temporal changes in cell and MV abundances, within the same order of magnitude (Figure 3A). Interestingly, the MV production rates during the early growth phase (0-24 hours) were 1-4 MVs cell^-1^ generation^-1^, with the exception of strain BS11 that exhibited a production rate of 64 ± 5 MVs cell^-1^ generation^-1^. During the late growth stages of the culture (24-72 hours) four out of six strains exhibited several-fold higher MV production rates (Figure 3B). The strain BS11 showed a strong decrease in the MV production rate, and in the strain HOT1A3 no changes were observed. We cannot exclude that higher abundances of MVs in “older” bacterial cultures could also represent MVs that were passively formed as a result of an increasing bacterial cell lysis (Toyofuku, Nomura and Eberl 2019). Nonetheless, a higher production of MVs at later stages of the bacterial growth could be a result of changing growth conditions in the culture (i.e., stress) (Orench-Rivera and Kuehn 2016). Furthermore, we speculate that the more active early stages of bacterial growth might have been associated with a higher turnover (i.e., higher uptake) of MVs, which potentially might have resulted in less MVs observed in media.

**Figure 3.**
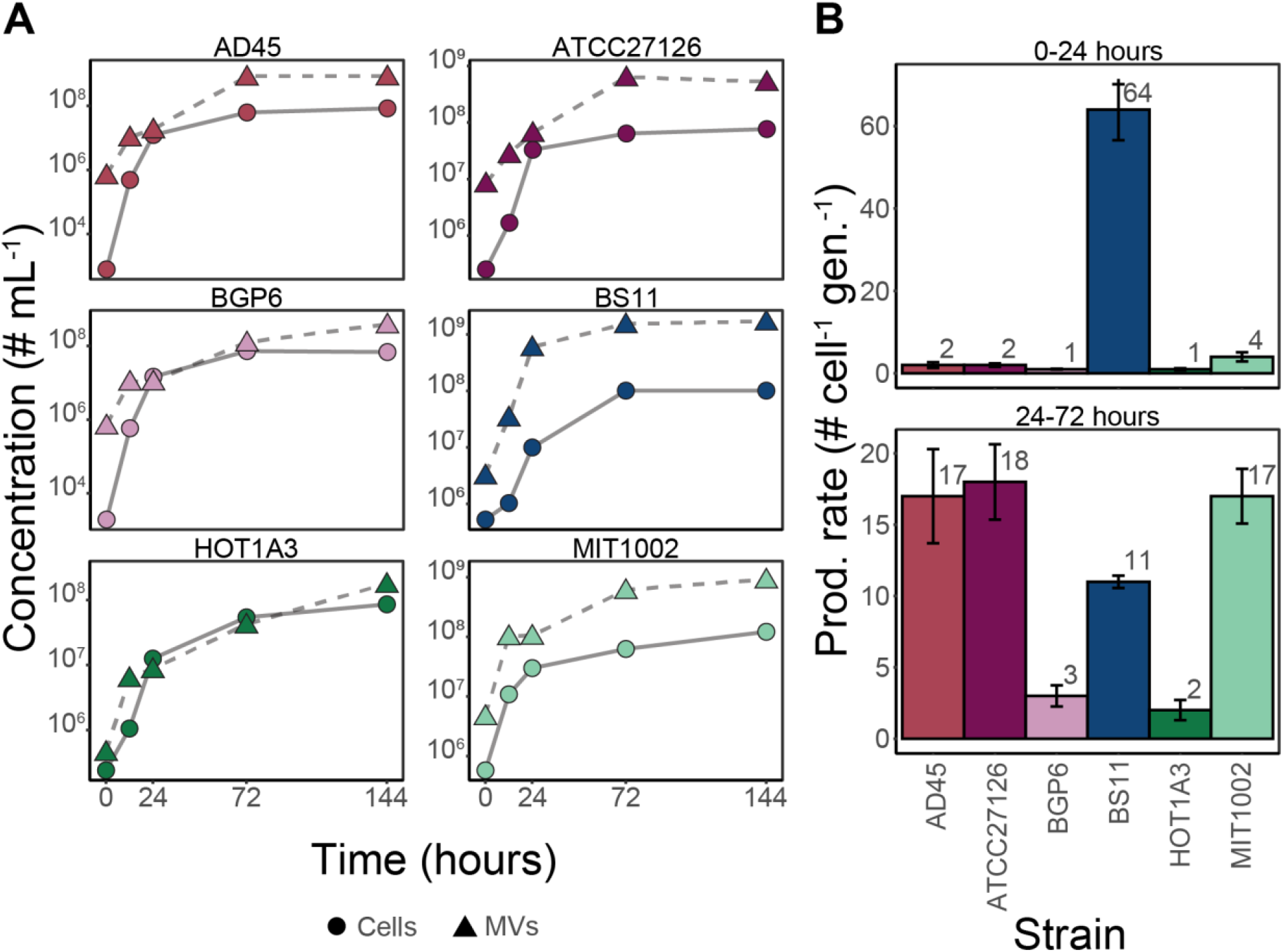
Production of MVs by *A. macleodii*. (**A**) Total cell (circles) and MV (triangles) abundances in six different strains of *A. macleodii*. The standard errors between the biological replicates are smaller than the symbols. (**B**) Calculated production rates of MVs during the early (0-24 h) and late (24-72 h) growth phases of each strain. The color coding of the individual strains is the same for panels A and B.

Although the size of MVs may affect their functionality (Turner *et al*. 2018), it is largely unknown whether there is a direct link between size and function (Nagakubo, Nomura and Toyofuku 2019). However, morphologically different MVs are likely mediating different biological functions (Toyofuku, Nomura and Eberl 2019). Due to the limited number of MVs that could be visualized by cryo-EM, we are not able to fully link the different morphologies and the quantitative size distributions. Nonetheless, in addition to outer- and outer inner-MVs, in some *A. macleodii* strains we also observed membrane-bound spherical and tube-shaped structures that enclosed large numbers of smaller MVs (Figure 1). These could potentially represent self-assembled structures, for example from lysed cells (Walker, Kennedy and Zasadzinski 1997; Turnbull *et al*. 2016). However, microscopic observations in various other bacteria suggest that such structures could also be produced by the bacterial cells and have specific functional roles (Kaplan *et al*. 2021). The origin of these structures in *A. macleodii* is unknown, however, morphologically similar structures in other marine bacteria were recently linked to sensing and utilization of polysaccharides (Fischer *et al*. 2019; Dürwald *et al*. 2021).

### The MVs of *A. macleodii* likely play a role in its copiotrophic lifestyle

To further elucidate the functional potential of MVs in *A. macleodii*, we characterized the protein content of the cellular (>0.22 μm) and the MV-associated fractions (100 kDa - 0.22 μm) in each strain. Overall, no significant differences in the proteome were found between the *A. macleodii* strains (PERMANOVA, p > 0.05) and more than 85% of all detected proteins in each strain were shared among all of them (Table 2). The consistency of the observations between the strains is a result of a high genomic similarity between them (average nucleotide identity > 90%; López-Pérez *et al*. 2012; Fadeev *et al*. 2016) and the similar non-limiting growth conditions applied here. Furthermore, the high portion of shared MV-associated proteins suggests potentially high intraspecific conservation of the produced MVs. Therefore, we focus below mainly on proteins observed in all the *A. macleodii* strains.

**Table 1.** *A. macleodii* strains used in this study.

| Strain | Origin | NCBI Reference Sequence | Reference |
| --- | --- | --- | --- |
| ATCC 27126<sup>T</sup> | North Pacific (surface) | NC_018632.1 | Baumann et al. 1984 |
| AD45 | Adriatic Sea (surface) | NC_018679.1 | Pujalte et al. 2003 |
| BS11 | Black Sea (surface) | NC_018692.1 | López-Pérez et al. 2012 |
| BGP6 | Pacific Ocean (surface) | NZ_CABDWP000000000.1 | Koch et al. 2020 |
| HOT1A3 | North Pacific (surface) | NZ_CP012202.1 | Fadeev et al. 2016 |
| MIT1002 | North Atlantic (surface) | NZ_CABDXM010000001 | Biller et al. 2015 |

**Table 2.** Overview of proteins detected in cellular and MV fractions of each *A. macleodii* strain.

| Strain | Total protein encoding genes | Proteins detected in cells | Proteins detected in MVs | Shared proteins* | Proteins detected only in cells* | Proteins detected only in MVs* |
| --- | --- | --- | --- | --- | --- | --- |
| **AD45** | 3817 | 1239 | 704 | 16 | 45 | 19 |
| **ATCC27126** | 3770 | 1512 | 1218 | 37 | 38 | 23 |
| **BGP6** | 3869 | 1356 | 1076 | 35 | 32 | 18 |
| **BS11** | 3677 | 1554 | 1317 | 45 | 32 | 20 |
| **HOT1A3** | 3839 | 1169 | 556 | 10 | 48 | 17 |
| **MIT1002** | 3845 | 1455 | 1279 | 31 | 41 | 32 |

\*Comparison of the top 5% of proportionally most abundant proteins

The MVs are generally fragments of the producing cell and thus, their associated proteins represent a subset of the cellular proteome (Schwechheimer and Kuehn 2015). From all the proteins encoded by each strain, 36 ± 5% were detected in the cellular and 27 ± 9% in the MV-associated fraction (Table 2). In the top 5% of the most abundant proteins in each strain more than half were fraction-specific (32-48 proteins in the cellular fraction and 17-32 proteins in the MV-associated fractions; Table S1). Additional proteins, found in both the cellular and the MV-associated fractions, had significantly different proportional abundance between the fractions (PERMANOVA, p < 0.05). Predicted localization of these most abundant proteins indicated in all six *A. macleodii* strains that the MV-associated fraction comprised a significantly higher portion of proteins with membrane- and periplasm-related origins than to the cellular fraction (Figure 4). The localization profiles of MV-associated proteins were overall in agreement with previous studies (Kulkarni, Swamy and Jagannadham 2014; Veith *et al*. 2015; Biller *et al*. 2021). The abundant proteins observed in the MV-associated fraction matched those found enriched in other bacterial MVs (Deatherage *et al*. 2009; Schwechheimer and Kuehn 2015). However, we also detected that a significant portion of the most abundant proteins in the MV-associated fraction had a cytoplasmic origin. Outer inner-MVs encapsulating the cytoplasmic matrix fraction were observed in the *A. macleodii* cultures (Figure 2). They likely contributed to the presence of cytoplasmic proteins in the MV-associated fraction (Pérez-Cruz *et al*. 2015). Nonetheless, based on the relatively high portion of proteins of cytoplasmic origins we cannot fully exclude some level of cellular contamination in the MV -associated fraction.

**Figure 4.**
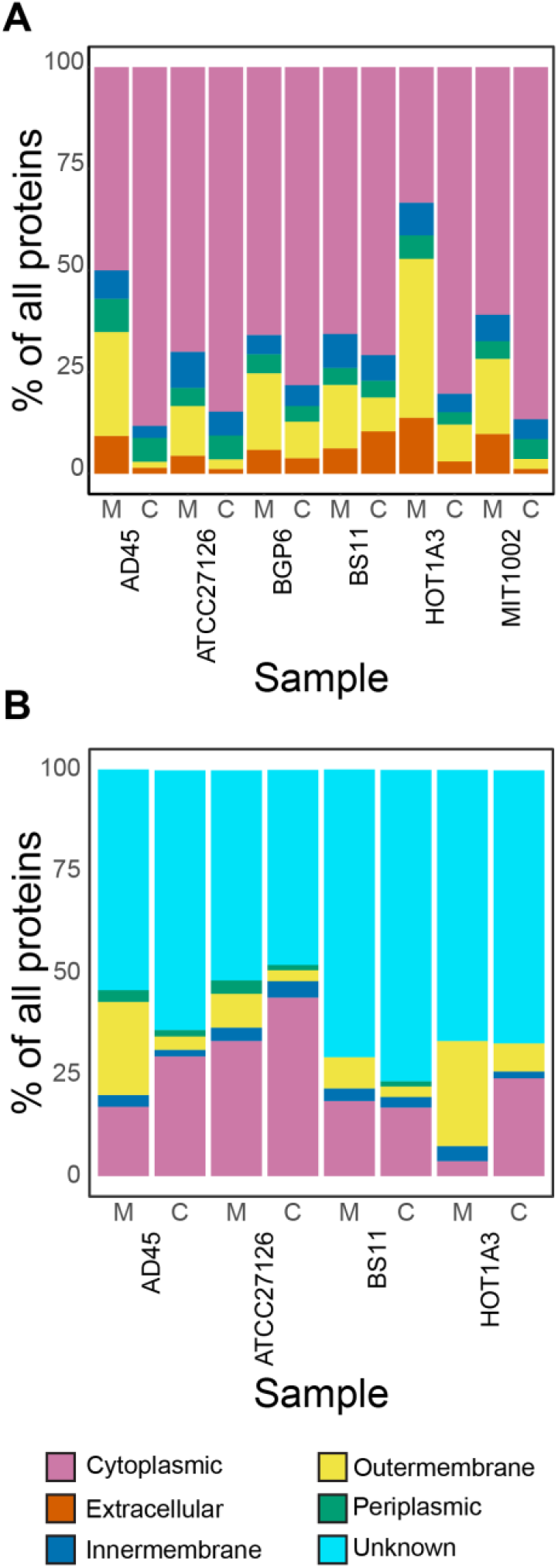
Proportions of subcellular localizations among the most abundant proteins in the cellular (‘C’) and MV (‘M’) fractions in each *A. macleodii* strain. The color code represents the different subcellular origins, predicted by CELLO2GO (**A**) and psortdb (**B**).

In all six *A. macleodii* strains the MV-associated fraction contained high proportion of various cell membrane related proteins (Figure 5). Compared to the cellular fraction, the MVs in all *A. macleodii* strains were characterized by a more than two-fold higher proportional abundance of outer-membrane proteins, such as *OmpA* and *OmpW*, inner-membrane type 1 secretion system transporters (*TolC*), and efflux channels such as the F0F1-type ATP synthase (both alpha and beta subunits) and the *MotA/TolQ/ExbB* proton channels (Table S1). However, the most prominent protein group was outer membrane *TonB*-dependent siderophore receptors reaching 7-33% of all detected proteins in MVs (Table S1). The TonB-dependent receptors mediate the transfer of various solutes such as organic molecules and siderophores via the outer membrane (Braun 1995). In *A. macleodii*, distinct sets of TonB-dependent receptors are expressed during carbon- and iron-limiting conditions (Manck *et al*. 2020) and during polysaccharide utilization (Neumann *et al*. 2015). Several non-marine gram-negative bacteria also produce MVs enriched in TonB-dependent receptors (Veith *et al*. 2009; Zakharzhevskaya *et al*. 2017; Dhurve *et al*. 2022). Although these receptors cannot transport solutes through the membrane, due to the absence of an energy source inside the MVs, they could still bind the solutes (Veith *et al*. 2015). Then, through fusion, bacterial cells could incorporate the MV-bound solutes such as siderophores (Prados-Rosales *et al*. 2014). In the ocean, such mechanisms of solute enrichment on MVs could potentially serve as community public goods that would support other bacteria such as lineages without iron-scavenging capabilities (Hogle *et al*. 2022; Manck *et al*. 2022).

**Figure 5.**
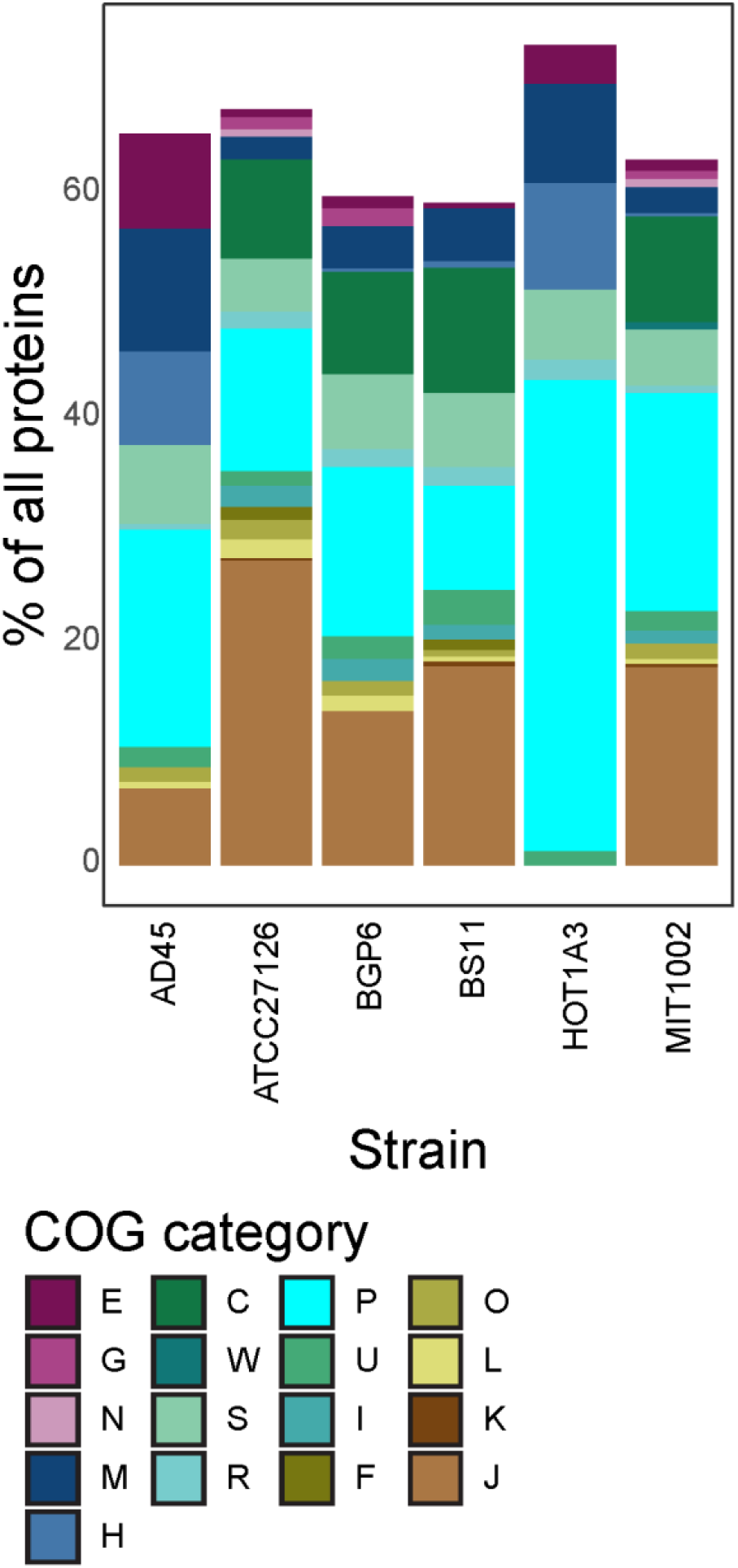
Proportion of selected COG categories among the most abundant MV-associated proteins in each *A. macleodii* strain. Included COG categories are: E - Amino acid transport and metabolism; G - Carbohydrate transport and metabolism; N - Cell motility; M - Cell wall/membrane/envelope biogenesis; H - Coenzyme transport and metabolism; C - Energy production and conversion; W - Extracellular structures; S - Function unknown; R - General function prediction only; P - Inorganic ion transport and metabolism; U - Intracellular trafficking, secretion, and vesicular transport; I - Lipid transport and metabolism; F - Nucleotide transport and metabolism; O - Posttranslational modification, protein turnover, chaperones; L - Replication, recombination and repair; K - Transcription; J - Translation, ribosomal structure and biogenesis.

Interestingly, in the MV-associated fraction of all six *A. macleodii* strains, we detected cytoplasmic proteins linked to cellular metabolic pathways, such as glutamine synthetase, superoxide dismutase, ketol-acid reductoisomerase, and riboflavin synthase (Figure 5). The MV-associated fraction also contained various proteins associated with post-translational modifications including chaperones (e.g., *GroEL*), thioredoxins (*TrxA* and *CnoX*), peptidylprolyl isomerase, and elongation factors (TufA, *Tsf*, and *FusA*; Table S1). Some of these proteins could potentially be associated with the cellular matrix encapsulated in outer inner-MVs (Kulkarni, Swamy and Jagannadham 2014; Park *et al*. 2015; Pérez-Cruz *et al*. 2015; Biller *et al*. 2021). However, others were moonlighting proteins with independent and different intra- vs. extracellular functionalities (Huberts and van der Klei 2010; Singh and Bhalla 2020). Such moonlighting proteins are present on membranes of gram-negative bacteria and their MVs (Park *et al*. 2015; Turnbull *et al*. 2016; Ebner and Götz 2019) and might suggest specific functional roles. For example, in *P. aeruginosa*, MVs enriched in various moonlighting proteins serve as a antibiotics resistance mechanism (Park *et al*. 2015). Among the proteins observed in this study, the chaperonin *GroEL*, responsible for proper protein folding inside the cytoplasm, can turn into a toxin outside of the cell (Yoshida *et al*. 2001). Additionally, glutamine synthetase, which was highly abundant in the MVs of all *A. macleodii* strains, can function as adhesion factor when exposed on the membrane (Candela *et al*. 2007). Similar adhesive functions have also been described for other proteins, such as the superoxide dismutase (Reddy and Suleman 2004) and the ketol-acid reductoisomerase (Castaldo *et al*. 2009). Based on observations on *Helicobacter pylori*, where MVs can enhance biofilm formation (Yonezawa *et al*. 2009), we tentatively assume that the potentially adhesive nature of MVs in *A. macleodii* facilitates its surface colonizing lifestyle (i.e., in biofilms; Ivars-Martinez *et al*. 2008).

It is important to note that, due to the considerable biomass required for the applied methodologies, we isolated *A. macleodii* MVs by targeting nanoparticles between 100 kDa - 0.22 μm. According to the minimal experimental requirements for defining EVs, the selected isolation strategy has an intermediate recovery and intermediate specificity (Théry *et al*. 2018). Therefore, based on existing knowledge regarding *A. macleodii*, it is likely that the selected size fraction also contained some other colloidal material (e.g., protein aggregates; López-Pérez *et al*. 2012). However, as such aggregates were not observed in microscopy and membrane-associated proteins were abundant, we are convinced that the targeted size fraction was highly enriched in MVs.

### The MVs harbor active hydrolytic ectoenzymes

The production of hydrolytic ectoenzymes is a prominent functional trait in marine heterotrophic bacteria and plays an important role in organic matter turnover (Arnosti 2011; Arnosti *et al*. 2021). The physiological benefit from releasing MV-associated ectoenzymes was demonstrated in *Escherichia coli*, where they strongly enhanced cellulose hydrolysis (Park *et al*. 2014). Among marine bacteria, active MV-associated hydrolytic enzymes were observed in *A. macleodii* KS62 isolated from seaweed (Naval and Chandra 2019), the coral pathogen *Vibrio shilonii* (Li, Azam and Zhang 2016) and *Prochlorococcus* (Biller *et al*. 2021). We found that the MVs of *A. macleodii* harbored a considerable amount and diversity of enzymes linked to bacterial extracellular enzymatic activity (EEA). Among these were leucyl aminopeptidases (EC 3.4.11.1), inorganic pyrophosphatases (EC 3.6.1.1) and putative alkaline phosphatases (EC 3.1.3.1). Other hydrolytic enzymes such as beta-glucosidase (EC 3.2.1.21) were also observed among MV-associated proteins although with low proportional abundance. The molecular weights of these ectoenzymes in *A. macleodii*, as characterized by the mass spectrometry analysis, were all below 100 kDa. Thus, unless associated with larger structures (i.e., MVs), they should have not been detected in the 100 kDa - 0.22 μm fraction. However, when dissolved, these enzymes can form polymeric complexes with high molecular weight (Barrett *et al*. 2012), which to some extent are retained in the MV-associated fraction.

Ectoenzymes in marine bacteria can be both outer-membrane bound and periplasmic, with higher activity measured in the latter (Martinez and Azam 1993). This is especially relevant for MVs that, as a result of their formation process, could function as enzymatic hotspots. To test whether the observed MV-associated ectoenzymes were active, we conducted biochemical assays on isolated MVs of the *A. macleodii* type strain ATCC27126. We targeted three different enzymatic groups commonly used to estimate marine bacterial extracellular enzymatic activity (EEA). Dependent on the enzymatic group, the total cell-free EEA of ATCC27126 represented up to 20% of the bulk enzymatic activity in the culture (Figure 6). Across all enzymatic groups, 60-90% of the cell-free EEA was associated with the MV fraction, with only a small portion of EEA measured in the < 100 kDa fraction (Figure 6). Previous size fractionated EEA measurements in natural seawater have found that up to 30% to the cell-free EEA was contributed by the MV-containing size fraction (0.02 - 0.2 μm; Baltar *et al*. 2019). Based on our results, we speculate that in *A. macleodii* nearly all the EEA is associated with MVs. These results encourage further targeted in-depth investigations of MV-associated enzymes in cultures of marine bacterial isolates as well as in natural seawater to estimate their contribution to the enzymatic turnover of organic matter in the ocean.

**Figure 6.**
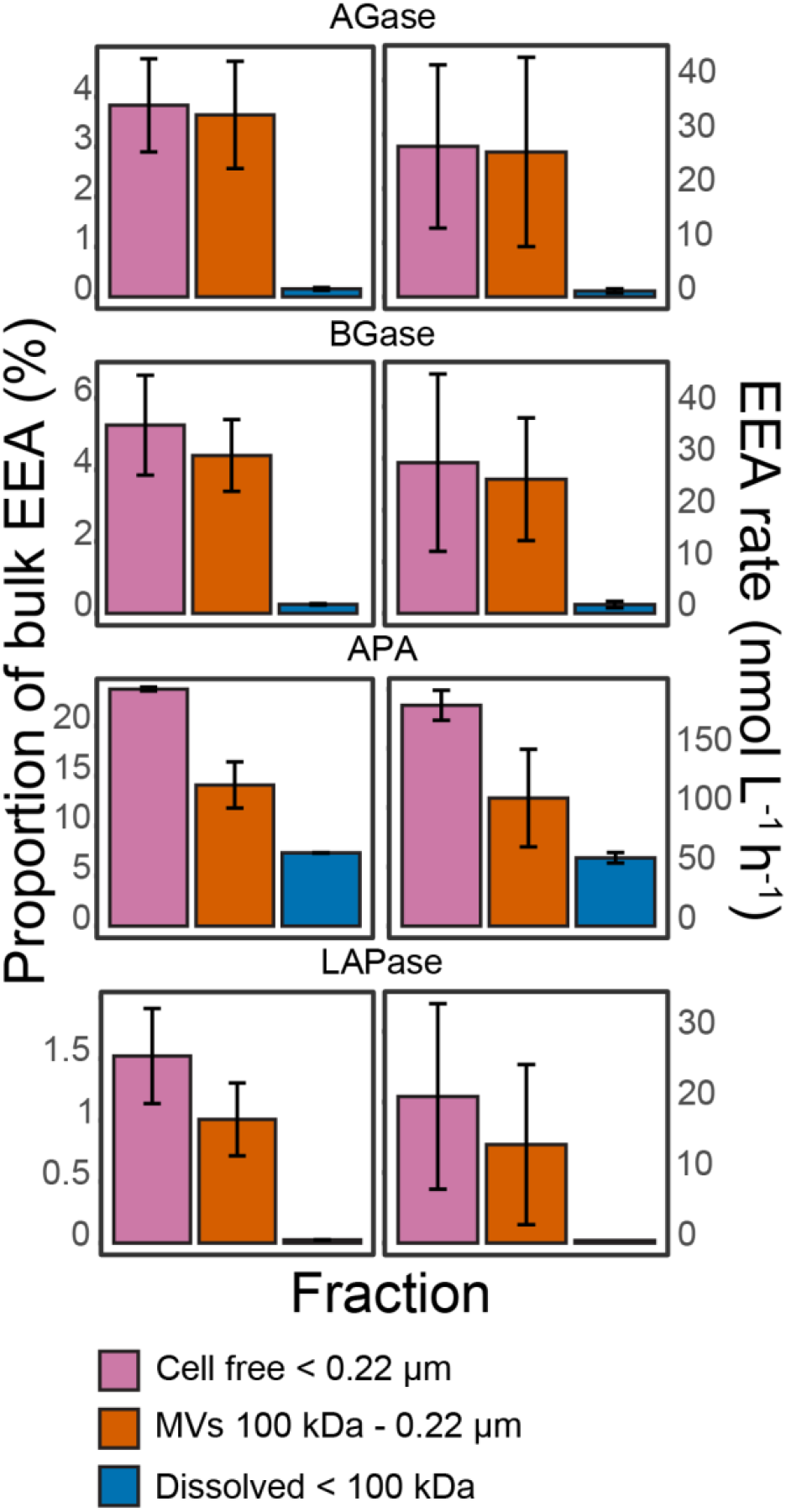
Extracellular enzymatic activity (EEA) of different size fractions in culture of *A. macleodii* type strain ATCC27126. The bulk represents EEA of non-filtered samples (i.e., bacterial cells + ectoenzymes). The color coding of the individual fractions is the same for all plots. AGase - alpha-glucosidase, BGase - beta-glucosidase, APA - alkaline phosphatase, LAPase - L-aminopeptidase.

## Conclusions

We showed that *A. macleodii*, a widespread and abundant heterotrophic marine bacterium, produces diverse MVs. Production and release of MVs are energetically costly and thus should have physiological and/or ecological benefits for the bacterial cells. While some of the observed characteristics correspond to findings in confined host-bacteria environments, the roles of MVs in marine bacterial assemblages remain largely unknown. In the oligotrophic marine environment extended scavenging capabilities are highly beneficial. Our results suggest that, through enrichment of receptors and adhesion proteins, released bacterial MVs could function as passive collectors of various solutes (e.g., iron and carbohydrates). The ability to produce and utilize such ‘hotspots’ of biologically relevant resources might provide an ecological advantage. Furthermore, in an environment in which heterotrophic cleavage of organic matter is conducted mostly by extracellular enzymes, the ability to secrete vehicles for hydrolytic enzymes might increase the efficiency of the extracellular enzymatic nutrient acquisition. Taken together, the observations in this study lay the ground for thorough, mechanistic investigations on MV diversity and function in the ocean with potentially important implications for the understanding of the oceanic microbiome and biogeochemistry.

## Material and Methods

### Culturing *A. macleodii* strains

Six *A. macleodii* strains (Table 1) were grown in artificial seawater PRO99 medium (Moore *et al*. 2007) amended with defined organic compounds (0.05% w/v of each lactate, pyruvate, acetate and glycerol) and a vitamin mix (Sher *et al*. 2011; Fadeev *et al*. 2016). Cultures were incubated at 20°C in the dark. To minimize sampling of cell debris they were not mixed during the incubation and before sampling.

To estimate MV production rates, each strain was grown in 12 biological replicates of 50 mL each. At every time point (after 12, 24, 72 and 144 h) three replicates from each strain were sampled. For protein analysis and morphological characterization of MVs, 2 L cultures of each *A. macleodii* strain were grown for 48 h. For EEA assays, *A. macleodii* ATCC27126 was grown in triplicate of 50 mL and sampled after 24 h. From each culture, 1 mL of cell suspension was preserved with 4% glutaraldehyde (final conc.) and stored at −80°C until flow cytometric analysis. The remaining culture was used for isolation of MVs.

### Bacterial cell enumeration

Bacterial cells were stained with 1× SYBR Green I (Thermo Fisher Scientific, MA, USA) for 15 min and then quantified using a flow cytometer (BD Accuri™ C6, BD, NJ, USA). The were processed using R package ‘flowWorkspace’ v4.8.0 (Finak and Jiang 2022) with hierarchical gating of singletons using R package ‘flowStats’ v4.8.2 (Hahne *et al*. 2022) and removal of cell debris using R package ‘openCyto’ v4.2 (Finak *et al*. 2014).

### Isolation of MVs

The MVs were isolated following the experimental guidelines for definition of EVs and their functions of the International Society for Extracellular Vesicles (Lötvall *et al*. 2014), and following Biller et al. (2022) for filtration size cut-offs for marine bacteria.

For production rates and EEA measurements, *A. macleodii* cultures were filtered manually using 50 mL syringes through a 25 mm diameter, 0.22 μm polycarbonate filter (Merck Millipore, MA, USA) to remove bacterial cells. To remove small molecular weight solutes, the filtrate was concentrated ca. 10 times using Amicon centrifugal filter units (Merck Millipore) with a 100 kDa cut-off to a final volume of 5 mL. The 100 kDa - 0.22 μm size fraction containing the MVs was then used for nanoparticle tracking analysis (NTA).

For protein analysis and morphological characterization, the 2 L cultures were filtered using a peristaltic pump through a 142 mm diameter, 0.22 μm polycarbonate filter (Merck Millipore, MA, USA) at a low pumping rate of 30 mL min^-1^ to minimize cell lysis. The filtrate was concentrated ca. 10 times using a tangential flow filtration with a 100 kDa cut-off (Vivaflow, Sartorius, Germany) to 200 mL, and further concentrated another ca. 10 times using Amicon centrifugal filter units (Merck Millipore) with a 100 kDa cut-off to a final volume of 20 mL. A 5 mL subsample of the concentrated MVs was collected for NTA. The remaining 15 mL of the concentrated MVs were pelleted using ultracentrifugation at 100,000×g at 4°C for 4 h and resuspended in 10 mL of clean PRO99 medium. After an additional round of ultracentrifugation, the pelleted MVs were resuspended in 250 μL of clean PRO99 medium. All isolated MVs samples (excluding those for EEA assays) were stored at −20°C until further analysis.

### Enumeration of MVs using nanoparticle tracking analysis

The abundance of MVs was measured by nanoparticle tracking analysis (NTA) using a NanoSight NS300 instrument (Malvern Panalytycal, UK) equipped with a 488 nm laser. Each MV sample was injected using a syringe pump with a flow speed of 100 μL min^-1^, and five videos of 60 sec were recorded. Videos were analyzed using NanoSight NTA software v3.2. For MV production rate measurements, the samples were analyzed with a consistent per-strain-threshold setting of 4-6. All particles in the size range of 20-450 nm were included in the final MV abundance estimates. To achieve high accuracy of the estimated MV size distributions, all concentrated MV samples contained 20-100 particles per frame following the manufacturer’s standard protocols.

The flow cell was thoroughly flushed with Milli-Q water (Merck Millipore) between samples and visually examined to exclude carry over. The performance of the instrument was routinely checked by measuring the concentration and size distribution of standardized 100 nm silica beads.

### MV production rate determination

The MV production rates were calculated following Biller *et al*. (2014) based on the assumption that each bacterial cell produces a constant number of MVs per generation (*r*). By estimating the number of generations based on changes in cell abundances and the change in MV concentrations, the MV production rate *r* can be calculated as follows:

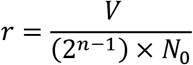

Where:

- *V* - number of produced MVs
- *N*_0_ - initial cell abundance
- *n* - number of generations

### Cryogenic electron microscopy (cryo-EM)

To verify the presence and size of MVs, 50 μL of the purified MVs was used for cryo-EM imaging. Plunge freezing and imaging was conducted at the Electron Microscopy Facility of the Vienna BioCenter on a Glacios transmission electron microscope (Thermo Fisher Scientific).

### Protein content characterization

For each *A. macleodii* strain, the protein content of cells retained on 0.22 μm filters and of 200 μL of the purified MVs (total of 10^10^-10^11^ MVs) were determined as described elsewhere (Bayer *et al*. 2019; Zhao, Baltar and Herndl 2020). Extracted proteins were subject to in-solution trypsin digestion and the resulting peptides were sequenced on a Q-Exactive Hybrid Quadrupole-Orbitrap Mass Spectrometer (Thermo Fisher Scientific) at the Vienna Research Platform for Metabolomics & Proteomics. Using the Proteome Discoverer v3.0 (Thermo Fisher Scientific), tandem mass spectrometry spectra were searched using SEQUEST-HT against the corresponding genome of each strain. Search parameters were: enzyme: trypsin, fragment mass tolerance: 0.6 Da, max. missed cleavages: 2, fixed modifications: carbamidomethyl (Cys), optional modifications: oxidation (Met). Percolator parameters were: max. delta Cn: 0.6, max. rank: 0, validation based on q-value, false discovery rate (calculated by automatic decoy searches) was 0.05. Protein quantification was conducted using the chromatographic peak area-based label-free quantitative method.

Protein coding genes from *A. macleodii* genomes were extracted using Anvi’o v7.0 (Eren *et al*. 2015) and annotated against the Clusters of Orthologous Groups of proteins (COG) database (Tatusov *et al*. 2000; Galperin *et al*. 2015). Further analysis was done in R v4.2.1 (R Core Team 2022) using RStudio v2022.02.3 (RStudio Team 2019) and R packages ‘phyloseq’ v1.40 (McMurdie and Holmes 2013) and ‘vegan’ v2.6-2 (Oksanen *et al*. 2022). Subcellular localization of the identified proteins was predicted using the CELLO2GO online tool (Yu *et al*. 2014) and supported by model predictions in PSORTdb v4.0 (Lau *et al*. 2021).

### Extracellular enzymatic activity (EEA) assays

The EEA assays were based on hydrolysis of fluorogenic substrate analogues that provide an estimate for leucine aminopeptidase (LAPase) activity, alkaline phosphatase (APA), alpha- and beta-glucosidase (AGase and BGase, respectively; Hoppe 1983). The substrate working solutions were prepared in 2- methaoxyethanol and kept at −20°C. Standard calibration curves were established with 7 to 15 different concentrations of the fluorophores MUF and MCA prepared in 0.22 μm filtered artificial seawater ranging from 1 nmol L^-1^ to 1 μmol L^-1^.

Three technical replicates (300 μL each) from the different size fractions (bulk, 100 kDa-0.22 μm, < 100 kDa) were incubated with the fluorogenic substrates in Greiner Bio-one 96-well non-binding microplates. Concentrations of each substrate were determined based on previously conducted enzyme kinetics (data not shown): 600 μmol L^-1^ of 4-methylcoumarinyl-7-amide (MCA)-L-leucine-7-amido-4-methylcoumarin (LAPase), 100 μmol L^-1^ methylumbelliferyl (MUF)-phosphate (APA), and 400 μmol L^-1^ for both MUF-α-D-glucopyranoside MUF-β-D-glucopyranoside (AGase and BGase, respectively). Incubations were carried out at room temperature in the dark. Fluorescence in each sample was measured after 1.5 h using a Fluorolog-3 fluorometer with a MicroMax 384 microwell plate reader (Horiba, Japan) at excitation/emission wavelengths of 365/445 nm and 380/440 nm for MUF and MCA, respectively. Culture media with substrate were used as blanks to determine background fluorescence, which was then subtracted from the sample fluorescence. Using the calibration curves for each fluorochromes, the linear increase in fluorescence over time was then transformed to enzymatic hydrolysis activity (nmol L^-1^ h^-1^).

## Data availability

The mass spectrometry proteomics data have been deposited to the ProteomeXchange Consortium via the PRIDE partner repository with the dataset identifier PXD037008.

All analytical workflows are publicly available on GitHub (https://github.com/edfadeev/Alteromonas_MVs/).

## Author contributions

EF - designed and conducted MV characterization experiments, analyzed the data, and wrote the manuscript under the guidance of GJH. CCB - conducted the enzymatic activity experiments and analyzed the results. JH - conducted NTA measurements of MV production. SJB, DS and MW - provided *A. macleodii* cultures and contributed to the interpretation of the results and the writing of the manuscript.

## Acknowledgments

The authors thank Mirjam Aubert for her assistance with the experimental work, as well as Dikla Aharonovich and Marlene Brandstetter for their assistance in producing and interpreting electron microscopy images of MVs. The authors also thank Susanne Erdmann for critical review of the manuscript.

This project was funded by the Austrian Science Fund (FWF) grant number M2797-B to EF, and grant number I4978-B to GJH.

